# Fucoidan-coated Layer-by-layer Lipid Nanoparticles for the generation of CAR-Macrophages

**DOI:** 10.64898/2026.07.01.735684

**Authors:** Victor Passos Gibson, Houda Tahiri, Samy Omri, Alessia Filippini, Justine Saber, Nancy Braverman, Mariane Cajubá de Britto Lira-Nogueira, Xavier Banquy, Pierre Hardy

## Abstract

Modulation of immune cells as therapeutic tools has gained significant clinical relevance in the treatment of cancer. Among them, macrophages represent a promising immunotherapeutic platform not only because they can internalize tumor material, but also because they profoundly shape the tumor microenvironment through cytokine production, antigen presentation, metabolic regulation, and modulation of other immune and stromal populations. Lipid Nanoparticles (LNPs) have enabled RNA therapies to the bedside and are thus considered the gold standard for gene delivery. However, optimizing LNPs for RNA delivery to macrophages remains an active area of investigation. Here, we propose the surface modification of unPEGylated LNPs using the Layer-by-Layer (LbL) approach for enhanced RNA delivery to macrophages. Specifically, we show that fucoidan, a sulfated polysaccharide, when at the outermost layer in the LbL process provides two physicochemical advantages to unPEGylated LNPs: (1) stability in PBS and (2) resistance to lyophilization in the presence of cryoprotectant. Additionally, fucoidan improves macrophage targeting and RNA transfection efficiency compared to previously synthesized hyaluronan-decorated LbL LNPs. Fucoidan LbL LNPs (Fuc-LNPs) preferentially accumulated in CD11b+ macrophages when co-cultured with U87 glioblastoma cells, which was not observed for control PEGylated LNPs. Furthermore, Fuc-LNPs induced a higher transfection of mRNA in primary human macrophages when compared to PEGylated control LNPs. Using the model mRNA encoding CAR@CD19, Fuc-LNPs generated CAR macrophages which mediated CD19 cell ablation in vitro. Altogether, these findings highlight the potential of the LbL strategy to modulate the targeting properties of LNPs, improving RNA delivery to human macrophages and encouraging further studies using LbL LNPs for the generation of CAR-Macrophages in the context of solid tumors.

## Experimental section

### Materials

Solvents and chemicals were obtained from Sigma (Oakville, Canada) and Thermo Scientific (Waltham, USA). Cholesterol, 1,2-dioleoyl-sn-glycero-3-phosphoethanolamine (DOPE), 1,2-dioleoyl-sn-glycero-3-phosphoethanolamine-N-(lissamine rhodamine B sulfonyl) (ammonium salt) (rhodamine-PE), and 1-oleoyl-2-12-[(7-nitro-2-1,3-benzoxadiazol-4-yl)amino]dodecanoyl}-sn-glycero-3-phosphoethanolamine (NBD-PE) were obtained from Avanti Polar Lipids (Alabaster, USA). 8-[(2-hydroxyethyl)[6-oxo-6-(undecyloxy)hexyl]amino]-octanoic acid, 1-octylnonyl ester (SM-102) was purchased from Cayman Chemical (Ann Arbor, USA). 1,2-Dimyristoyl-sn-glycero-3-methoxypolyethylene glycol-2000 (DMG-PEG_2k_) was purchased from Nanosoft Polymers (Winston-Salem, USA). (DiIC_18_(7); 1,1′-dioctadecyl-3,3,3′,3′-tetramethylindotricarbocyanine iodide) (DIR) and aminooxy-Cyanine647 were purchased from Biotium (Fremont, USA). Fucoidan from *Fucus vesiculosus*, Poly-L-Arginine (PLR, 5-15 kDa), Sodium Periodate, 6-(*p*-Toluidino)-2-naphthalenesulfonic acid sodium salt (TNS), and Pur-A-Lyzer mini dialysis kit (6-8 kDa) were obtained from Sigma (Oakville, Canada). Hyaluronic acid (HA) (60 kDa) was obtained from Lifecore Biomedical Inc. (Chaska, USA). Oligonucleotides (siRNA and mRNA) were obtained from Horizon Discovery (Waterbeach UK), VectorBuilder (Chicago, USA), APExBio (Houston, USA), and Helix Biotech (Knoxville, USA). Sequences, whenever available, are provided in Table 1.

**Table 1.**
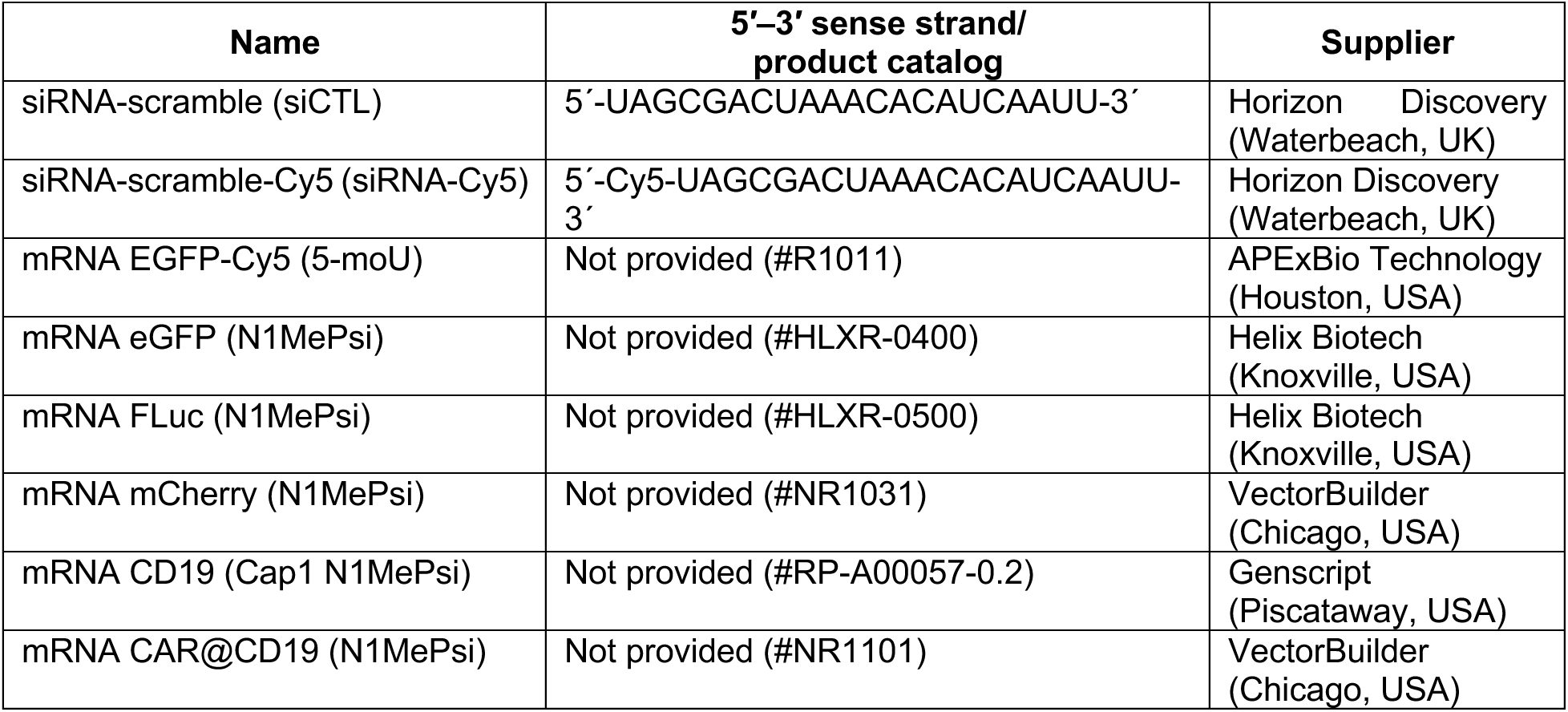
Sequences and suppliers of siRNA and mRNA.

### Synthesis of Layer-by-Layer LNPs and LNPs-PEG

Layer-by-Layer LNPs were synthesized as previously described (1) with slight modifications. Briefly, LNPs were synthesized by ethanolic injection where 45 μL of sodium acetate (10 mM, pH 4) solution containing mRNA (40:1 total lipid:mRNA w/w) is rapidly added into a 15 μL lipidic solution (0.22 μmol total lipids) containing SM-102:Chol:DOPE (LbL LNPs, 50:39:11 % mol) or SM-102:Chol:DSPC/DOPE:DMG-PEG_2k_ (LNPs-PEG, 20:38.5:10:1.5 % mol). The suspension was allowed to stand for 15 min at room temperature and 5 x diluted in sodium acetate. Layer-by-Layer coating was constructed by incubating 14.1 μL of unPEGylated LNPs with 7 μL of siRNA (20 μM, sodium acetate buffer), 1 μL of PLR (2 mg mL^-1^, nuclease-free water), and 5 μL of either hyaluronic acid (5 mg mL^-1^ in nuclease-free water) or fucoidan (1. 25 mg mL^-1^) to obtain HA-LNPs or Fuc-LNPs. Whenever applicable, LbL LNPs were dialyzed against PBS at room temperature for 2h for ethanol removal and raise the pH. For CAR experiments, the LNPs were synthesized with the double lipid molarity (0.44 μmol). The LbL process for 2x LNPs was the following: 14.1 μL LNPs, 12 μL of siRNA, 2 μL of PLR, and 5 μL of fucoidan (4 mg mL-1). No physicochemical differences were observed between 1x or 2x Fuc-LNPs.

### Physiochemical characterization of nanoparticles

Dynamic light scattering (DLS) and ζ-potential measurements were carried out on LiteSizer DLS 700 (Anton Paar Canada Inc, Montreal). Size and polydispersity Index (PDI) were measured with a 1 mm path length disposable cuvette, side scattering angle (90 °), 25 °C, 1 minute equilibration time, refractive index of 1.45, for 60 runs; ζ-potential runs were done using Anton Paar omega cuvette. Encapsulation efficiency was carried out using RiboGreen reagent diluting particles either 40 or 80 x in buffer containing or not 1% triton. For LbL LNPs, particles were diluted 40x, incubated with RiboGreen reagent, and total fluorescence was compared to equimolar RNA solution containing a mix of mRNA and siRNA. Additionally, LbL LNPs were formulated with either mRNA-Cy5 or siRNA-Cy5, prepared as usual, diluted to reach at least 10^9^ particles mL^-1^, and run through Nanoparticle flow cytometry (NanoFCM) to determine the % of loaded nanoparticles. Colocalization on NanoFCM were carried out with Cyanine647-aminooxy Fucoidan and unPEGylated LNPs core doped with NBD-PE (0.1% mol), detected at lasers 638 and 488 nm, respectively, with events between 3,000 and 12,000. For SAXS measurements, samples were concentrated two-fold using Amicon filter units (100 kDa, Thermo Scientific). Analyses were carried out at Structural Biology Platform of the Biochemistry department, Université de Montréal, using a BioXolver SAXS system (Xenocs) equipped with a MetalJet D2 + 70 kV X-ray source (Excillum) and a PILATUS3 R 300K detector (Dectris), after 30 s exposure. Average scattering profiles were obtained and subtracted from matching buffer scattering.

### Cell culture

HEK-293 was purchased from ATCC (Manassas, USA) and maintained in Dulbecco’s modified Eagle’s medium (DMEM; Thermo Scientific, Waltham, USA) supplemented with 10% FBS (Wisent, Canada) and 1% PenStrep (Wisent, Canada). Human monocytes (THP-1) were acquired from ATCC and cultured in RPMI-1640 (Wisent) supplemented with 10% FBS, 1 % PenStrep, and 2-Mercaptoethanol (0.05 mM, Thermo Fisher Scientific). Raji-GFP/Luc cells were purchased from Vitro Biotech (Austin, USA) and maintained similarly to THP-1 cells. U87-MG-GFP/Luc was purchased from Vitro Biotech (Austin, USA) and maintained in minimum essential medium (MEM, Thermo Scientific) supplemented with 1% non-essential amino acids (Gibco), 1% sodium pyruvate (Gibco), and 1% PenStrep or RPMI-1640 supplemented with 10% FBS and 1% PenStrep, respectively. THP-1 cells were differentiated in macrophages after 48h incubation with phorbol 12-myristate 13-acetate (10 ng mL^-1^, PMA), when media was renewed, and cells rested overnight. Primary monocytes were obtained from whole blood samples obtained from healthy individuals with authorization of the Research Ethics Board from the University Hospital CHU Sainte Justine (# 2026-8512). Monocytes were isolated by magnetic negative selection (EasySep Direct Human Monocyte Isolation Kit, StemCell Technologies, Vancouver, Canada) and cultured for 6 days in ImmunoCult^TM^-SF Macrophage Medium (StemCell) supplemented with 50 ng mL^-1^ of Human Recombinant M-CSF (StemCell) at the day of seeding and again on day 4. For downstream processing, human macrophages were detached with Accutase (Biolegend, San Diego, USA) or Trypsin/EDTA (Wisent) for 15 minutes at 37 °C.

### Co-culture experiments

Human monocytes (THP-1) were differentiated into macrophages for 48h after which the media was removed, cells washed in PBS, and U87-GFP-Luc cells were seeded and attached overnight. Cells were grown in U87 media (no effect on THP-1 cellular viability effect was observed). The next day, fluorescently labelled nanoparticles (DiR, FACS, Rhodamine-PE, microscopy, 0.1% mol ratio) were incubated for 1h, cells were washed and detached with Accutase. For labelling, Fc receptors were blocked (Human TruStain FcX) and macrophages were identified using PE anti-human CD11b antibody (#301305, Biolegend) according to manufacturer’s protocol. Cells were washed thrice and resuspended in facs buffer. For microscopy experiments, cells were culture as described, but int he presence of a coverslip. Cells were treated, fixed with 4% PFA solution in PBS, washed, stained with Hoechst 33342 (Cayman Chemical) mounted on a coverslip and imaged using Laser scanning confocal microscope with spectral emission detection (Leica TCS SP8, Wetzlar, Germany).

### Cellular Transfection

For transfection experiments, LbL LNPs or PEG-LNPs were diluted to reach the desired concentration indicated in the legend of each experiment and incubated for 48h unless stated otherwise. For eGFP and mCherry expression, cells were detached with Trypsin/EDTA for 5 or 15 minutes (macrophages only), neutralized with full media, washed with PBS thrice, and resuspended in Facs Buffer (PBS supplemented with 2 % FBS) and acquired on BD LSRFortessa. For FLuc experiments, cells were lysed directly on the well-plate following the Steady-Glo reagent protocol (Promega, Madison, USA) and luminescence was measured on a CLARIOstar plate reader (BMG Labtech, France). Cellular viability was carried out with PrestoBlue reagent (Thermo Scientific). For the inhibition assay, 10 k monocytes were differentiated into M0 macrophages in a 96-well plate as previously described and incubated with either 50 or 100 μg mL^-1^ of fucoidan for 1h, when fucoidan-containing media was renewed, and rhodamine-labelled nanoparticles were added for 2h. After the incubation time, media was removed, cells were washed thrice in cold PBS, and lysed with 100 μL of the disruption solution containing 0.5 mg mL^-1^ heparin (in PBS) and DMSO (1:1 v/v ratio) (2).

### CAR-Macrophages

To generate CAR-Macrophages, THP-1 or human primary derived macrophages were differentiated as described and incubated with Fuc-LNPs or PEG-LNPs for 48h, cells were washed with PBS and processed for CAR detection by flow cytometry or incubation with Raji cells. To detect the presence of CAR@CD19, macrophages were detached with Accutase, blocked with Human TruStain FcX, and incubated with FITC-Labeled Human CD19 (CD9-HF2H2, ACRO Biosystems, Newark, USA) according to manufacturer’s instructions. Cells were washed thrice and resuspended in facs buffer (PBS supplemented with 2% FBS). For cytotoxicity essays, CAR-Macrophages were detached with Accutase, washed, counted, and incubated for 48h with 20 k Raji-GFP-Luc cells at different effector-to-target ratios, varying only the number of CAR or untransfected macrophages. Co-incubation with U87-GFP/Luc expressing CD19 cells followed similar protocol, but U87 cells were transfected 24h prior co-incubation with mRNA encoding CD19 protein (GenScript, NJ, USA) using Lipofectamine 3000 (Thermo Scientific) following the manufacturer’s protocol. The expression of CD19 was detected by flow cytometry using RR688 anti-huCD19 (#572375, BD Biosciences, San Jose, USA). 24h after co-culture, supernatant was removed for the detection of TNF-α cytokine (#430207, BIolegend). The expression of Signal regulatory protein α (SIRPα, APC anti-hu CD172a, #372105, Biolegend)) and CD47 (APC anti-hu CD47, #323123, Biolegend) was detected by flow cytometry 24h after seeding. Live images were acquired on the Incucyte S3 Live Cell Analysis System (Sartorius, Germany) and cells were lysed directly on the well-plate using Steady-Glo reagent to measure the luciferase activity corresponding to Raji cells. For confocal images, untransfected or CAR-M were co-incubated with U87-GFP/Luc cells expressing CD19 in a 24-well plate containing a coverslip. After 24h, cells were wash thrice in ice-cold PBS, fixed in a 4% paraformaldehyde in PBS for 15 minutes at room temperature, followed by two additional PBS washes. Fixed samples were permeabilized and blocked in 0.1% Triton X-100 and 10% FBS in PBS for 1 h, then incubated overnight at 4 °C with rabbit anti-human IBA1 (1:400, Wako 019-19741). The next day, coverslips were washed and incubated with Alexa Fluor 594-conjugated anti-rabbit secondary antibody (1:450, Invitrogen 21207). Nuclei were counterstained with DAPI, and coverslips were mounted for imaging. Confocal images were acquired on a Zeiss LSM780 laser-scanning microscope with DIC collected in parallel to visualize cell morphology.

### Statistical analysis

Experiments were performed in triplicate unless stated otherwise and expressed as mean ± standard deviation. Data were plotted in Prism Software (GraphPad Software 10, La Jolla, CA, USA), and statistical significance was calculated with two-tailed Student’s t-test, one-way ANOVA with post hoc comparison through Tukey’s test, or two-way ANOVA with Sidak ’s multiple comparison, and expressed with their respective p values.

## Introduction

A great deal of research has been focused on understanding and targeting non-cancerous cells during tumorigenic process (3, 4). Within the tumor microenvironment, macrophages play a key role in tumour progression and maintenance (5, 6), fostering the development of immunotherapeutics able to modulate their fate by mediating their depletion (7, 8), re-education or reprogramming (9), or enhancing their effector functions (10). In this investigation, we concentrate on the latter by harnessing the therapeutic potential of nanomedicine (11).

Lipid nanoparticles (LNPs) represent a promising delivery platform in nanomedicine due to their biocompatibility, low toxicity, and ability to encapsulate a wide range of therapeutic molecules. These properties have led to their increasing use, particularly in mRNA delivery, as demonstrated by the development of COVID-19 vaccines (12). The LbL strategy is a well-known methodology employed in material sciences to modulate the properties of surfaces through the deposition of alternating layers of opposingly charged polyelectrolytes. It has been largely used to modify lipidic and polymeric-based nanomaterials (13), leading to profound biodistribution (14, 15) and therapeutic effects (16). We have previously employed the LbL approach to modify unPEGylated LNPs containing RNA interference. By leveraging hyaluronic acid as the outermost layer, we promoted glioblastoma targeting of nanoparticles enhancing the antitumor effect of microRNA-181a-5p both *in vitro* and *in vivo* (17, 18). Here, we sought to expand the applicability of LbL LNPs by redirecting the system to macrophages harnessing their therapeutic benefits through messenger RNA modulation.

Macrophages are central effectors of innate immunity, endowed with functional plasticity that allows them to participate in both inflammatory processes and tissue resolution. Their pivotal role in phagocytosis, antigen presentation, and modulation of the tumor microenvironment makes them strategic candidates for anticancer immunotherapies (5). In this context, the engineering of CAR-macrophages (CAR-M) has emerged as an innovative approach. These cells are genetically modified to express a *Chimeric Antigen Receptor* (CAR), granting them the ability to specifically target tumor cells. While CAR-T cells have demonstrated remarkable success in hematological malignancies, their efficacy against solid tumors remains limited due to physical barriers, immunosuppressive signaling, and antigen heterogeneity. Unlike CAR-T cells, CAR-Ms exploits their natural ability for tissue infiltration and immune remodeling to induce a more robust antitumor response (19). This strategy, still in preclinical and early clinical development (20, 21), represents a promising advancement in the field of cellular immunotherapy.

Fucoidan is a sulfated polysaccharide primarily derived from brown seaweed. Its bioactive properties include antioxidant, anti-inflammatory, antitumor, and immunomodulatory effects, which have been extensively investigated in various preclinical models (22–25).

In addition, the strong affinity of fucoidan for macrophage and related receptors, as P-selectin, facilitates selective nanoparticle uptake (26, 27). As a negatively charged polysaccharide, fucoidan is an ideal potential candidate for the LbL process. Therefore, in this investigation, we constructed fucoidan-terminated LbL LNPs (Fuc-LNPs) using the LbL assembly to generate CAR-macrophages. By using fucoidan as the outermost layer, we bestowed colloidal stability to mRNA-loaded unPEGylated LNPs in iso-osmolar media (PBS) in additional to resistance to lyophilization. Finally, using several reporter and functional mRNA templates, we confirmed Fuc-LNPs successfully transfected immortalized and primary human-derived macrophages, generating active CAR-Ms active against CD19-expressing tumour cells.

## Results

### Synthesis and characterization of LbL LNPs

Starting with unPEGylated LNPs composed of SM102:Cholesterol:DOPE (50:39:11 mol %), we proceeded with the layer-by-layer coating by firstly layering the lipidic template with scramble siRNA, then Poly-L-Argine (PLR), and either Hyaluronic Acid (HA) or Fucoidan (Fuc) to synthesize HA-LNPs and Fuc-LNPs, respectively (Figure 1A). Physicochemical properties during each step of LbL synthesis are described in Figure 1B. A gradual increase in particle size was observed and albeit the slight increase in the polydispersity Index (PDI), values remained below 0.3, indicating little aggregation during the LbL assembly. Finally, we observed an inversion of the ζ-potential after each layering step as it is expected during the layer-by-layer coating of nanoparticles. Initial unPEGylated LNPs core were synthesized with size of 148 ± 3 nm, PDI of 0.127 ± 0.01 and ζ-potential of 59 ± 1 mV. Final HA-LNPs presented size, PDI, and ζ-potential of 192 ± 9 nm, 0.158 ± 0.05, and -22 ± 1, respectively, and Fuc-LNPs were obtained with generally similar size and PDI (182 ± 6 nm, 0.143 ± 0.01), but higher modular ζ-potential (−39 ± 1 mV).

**Figure 1.**
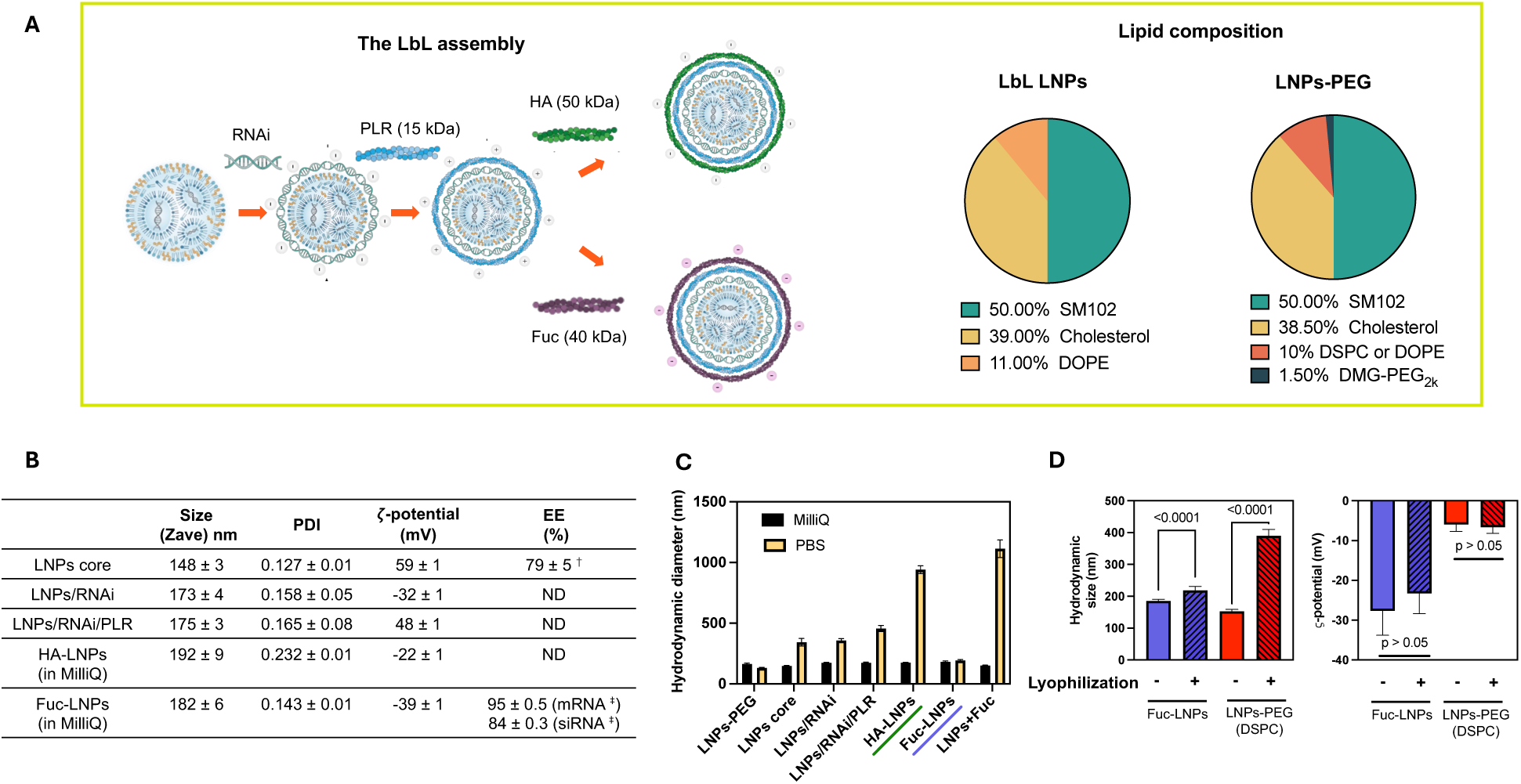
Physicochemical characterization of LbL LNPs. (A) Schematic representation LbL LNPs terminated with hyaluronic acid (HA-LNPs) or fucoidan (Fuc-LNPs). The LNPs core (SM02:Chol:DOPE 50:39:11 mol%) is first layered with siRNA, then Poly-L-Arginine (PLR), and finally sodium hyaluronate or fucoidan. (B) HA-LNPs and Fuc-LNPs characterized by dynamic light scattering (DLS) and electrophoretic mobility. Encapsulation efficiency of mRNA unPEGylated core was determined by RiboGreen assay (^†^), whereas mRNA and siRNA loading after LbL assembly was determined by Nano flow cytometry (^‡^). (C) At each step of the LbL process, nanoparticles were diluted in pure water or PBS. (D) Fuc-LNPs and LNPs-PEG (DSPC) were formulated per protocol, diluted (1:1 v) in sucrose 10% and lyophilized for 48h. Formulations were resuspended in PBS and characterized by hydrodynamic size (reported as z-average), PDI, and ζ-potential.

As expected, due to their unPEGylated nature, the lipidic core, as well as LNPs after the first and second LbL step (LNPs/RNAi and LNPs/RNAi/PLR, respectively) were not stable upon dilution in PBS, as the neutralization of ionizable lipids and the absence of a hydrophilic anchored lipid (*e.g.* DMG-PEG_2k_) drives colloidal instability at pH 7.4 (Figure 1C). Hyaluronan-decorated LbL LNPs (HA-LNPs) were also not stable in PBS, but, surprisingly, fucoidan at the outermost layer protected unPEGylated LNPs from PBS destabilization. Regarding particle size, the hydrodynamic diameter of HA-LNPs after dialysis against PBS raised from 175 ± 3 nm to 941 ± 32 nm, whereas Fuc-LNPs remained stable with size varying from 182 ± 7 to 193 ± 7 nm (Figure 1C). However, HA-LNPs were stable upon dilution in PBS doped with 1% FBS, suggesting that protein corona formation is important for the colloidal stability of hyaluronan-decorated LbL LNPs (Figure S1A).

To better understand this observation, we coated the unPEGylated LNPs core directly with fucoidan bypassing the RNAi and PLR layers. We then dialyzed the fucoidan-layered LNPs against PBS. The fucoidan coated LNPs (LNPs + Fuc) with initial size of 151 ± 3 nm, PDI 0.13 ± 0.03, and ζ-potential -35 ± 2 mV, destabilized to 1114 nm and PDI > 0.3, but remained negatively charged (−50 ± 1 mV) after dialysis (Figure 1C, LNPs + Fuc). We also confirmed the PBS stability of the three-layer Fuc-LNPs by Nano Flow Cytometry (NanoFCM) where again no significant changes in particle size were observed when particles were diluted in pure water or PBS. However, NanoFCM revealed a slight decrease in nanoparticles’ concentration in PBS (1.17 ± 0.21E^10^ NPs mL^-1^ v. 1.47 ± 0.22E^10^ in pure water) accompanied by a right shift in size distribution (Figure S1B).

Collectively, these findings indicate that the final layer-by-layer construction bearing fucoidan as outermost layer, and not hyaluronic acid, is essential for the stability in PBS of layer-by-layer coated mRNA-loaded unPEGylated LNPs. Interestingly, the complete three-layered fucoidan-decorated LbL LNPs is crucial for colloidal stability in PBS as intermediate LbL species and directly layered fucoidan unPEGylated LNPs were not.

After observing that fucoidan provided colloidal stability to LbL LNPs in a iso osmotic pH 7.4 solution, we wondered if these nanoparticles could also withstand lyophilization, an important hurdle in LNPs research. We formulated mRNA luciferase-loaded Fuc-LNPs as per protocol and mixed nanoparticles (1:1 v/v ratio) with a 10wt% sucrose cryoprotectant solution. After lyophilization, Fuc-LNPs were resuspended in PBS. PEGylated LNPs were formulated using Moderna Inc. vaccine formulation (28), henceforth named LNPs-PEG (DSPC), for comparison purposes (Figure 1A). Remarkably, Fuc-LNPs were stable after lyophilization as little variation in particles’ size was observed, which was not the case for PEGylated LNPs (Figure 1D). Before lyophilization, Fuc-LNPs mixed with cryoprotectant presented size, PDI, and ζ-potential of 185 ± 5 nm, 0.15 ± 0.07, -28 ± 6 mV, respectively. Right after lyophilization, nanoparticles were resuspended and physicochemical parameters changed to 218 ± 12, 0.26 ± 0.03, -23 ± 5. On the other hand, PEGylated LNPs mixed with cryoprotectant increased in size from 153 ± 6 before lyophilization to 390 ± 19 nm after lyophilization (Figure 1D). Importantly, Fuc-LNPs induce similar RNA transfection in HEK 293T cells *in vitro* than freshly synthesized Fuc-LNPs (Figure S1C). These findings suggests that fucoidan not only provide colloidal stability to unPEGylated LNPs when at the outermost layer in the LbL process but also enable the lyophilization of these particles while maintaining efficient mRNA transfection *in vitro*.

Taking advantage of the nano flow cytometry platform, we addressed two challenges when developing LbL LNPs constructs: (1) determining the encapsulation efficiency of both core and layered genetic cargo after LbL assembly (Figure 1B and Figure S2A) and (2) providing evidence to the integrity of the three-layered system synthesized herein (Figure S2B). By encapsulating Cyanine 5-tagged mRNA or siRNA, we determined the % of RNA-loaded particles at the end of the LbL process (Figure 1B). For mRNA, the calculated encapsulation efficiency was 95 ± 0.5 % for Fuc-LNPs. Comparatively, the mRNA encapsulation efficiency measured by RiboGreen fluorescence assay, which is done exclusively in the unPEGylated core template, reached 79 ± 5 %. The encapsulation efficiency for the RNAi layer reached 84 ± 0.3% as per nanoFCM analysis.

The nano flow cytometry platform allows us to distinguish RNA-loaded from empty nanoparticles by tracking the positive and negative Cy5 population, respectively (Figure S2A for representative images). The % for unloaded and loaded populations at the end of the LbL process reached 61 ± 1 v. 39 ± 1 % when tracking cyanine 5-tagged mRNA, and higher than 99% for loaded nanoparticles when tracking cyanine 5-tagged siRNA.

Finally, driven to provide more information on the structure of LbL LNPs, we formulated fluorescently labelled unPEGylated LNPs by incorporating 18:1-12:0 NBD PE in the lipidic composition. Tagged particles were then used as template to construct Fuc-LNPs and a fluorescent Fuc-LNPs where the outermost layer was assembled using Cyanine 647-tagged fucoidan. The NBD+ population accounted for more than 90% of total Fuc-LNPs, confirming the efficient incorporation of the tagged lipid in the final Fuc-LNPs construct (Figure S2B, left graph). In the dual NBD and Fuc647 labelled system, the double positive population accounted for 89 ± 3 % of Fuc-LNPs, within the NBD+ population, suggesting the main system is composed by a structurally organized nanoparticle containing a LNP core and a Fucoidan layer (Figure S2B, right graph).

We then addressed if the enhanced colloidal stability of Fuc-LNPs originated from a shielding effect provided by the fucoidan-terminated three-layered structure. We first tested the accessibility of the ionizable core within the Fuc-LNP construct using the 6-(p-toluidino)-2-naphthalenesulfonic acid sodium salt (TNS) assay which revealed a slight increase in the apparent pKa of Fuc-LNPs (pK_a_^A^ 6.72) when compared to their PEGylated counterparts (pK_a_^A^ of 6.72 and 6.36, DSPC and DOPE as helper lipid, respectively) (Figure S3A). Secondly, we used the RiboGreen dye assay to assess the RNA content in the LbL assemblies comparing hyaluronan v. fucoidan terminated LbL LNPs. Compared to a free equimolar RNA solution, both LbL systems demonstrated cargo protection, with Fuc-LNPs presenting lower fluorescence signal than the HA counterpart (Figure S3B).

Finally, we submitted Fuc-LNPs and PEGylated LNPs to Small Angle X-ray Scattering (SAXS) to identify changes in the morphological structures of nanoparticles upon dilution in either pure water or PBS. For the SAXS analysis, noticeable peaks were observed for PEGylated LNPs containing either DOPE and DSPC, unPEGylated LNPs core, and Fuc-LNPs upon dilution in pH 4 sodium acetate buffer (10 mM). PEGylated and unPEGylated LNPs shared similar bragg’s diffraction peaks at q_1_ = 0.139 Å^-1^, but Fuc-LNPs presented a more intense left-shifted peak (q_2_ = 0.126 Å^-1^), which disappeared upon dilution in PBS, whereas PEGylated LNPs lost resolution but still presented a faint left-shifted peak (q_1_ = 0.131 Å^-1^) compared to their water-diluted counterparts. We did not perform SAXS experiments for unPEGylated LNPs at pH 7 due to colloidal instability in this solution (Figure S3C).

### Fuc-LNPs mediates mRNA transfection in vitro and in vivo

After synthesizing and characterizing LbL LNPs, we then investigated the cellular interaction of both Fuc-LNPs and HA-LNPs with human-derived macrophages (HDM) differentiated from immortalized human THP-1 cells. To initially attest if sulfated polysaccharide fucoidan confers LbL LNPs a tropism towards macrophages (M0), we first studied the uptake of HA-LNPs and Fuc-LNPs by M0 cells (Figure 2A). The uptake experiments were conducted in full media (RPMI supplemented with 10% FBS) to ensure the colloidal stability of HA-LNPs. As expected, Fuc-LNPs demonstrated superior binding to M0 cells at the three time points tested. As early as 15 minutes, the uptake of M0 with DIR-labelled Fuc-LNPs reached 65% of total population (v. 35% for HA-LNPs, Figure 2A). After 30 and 60 minutes, the uptake of Fuc-LNPs was 73 and 85%, respectively (v. 53 and 77% for HA-LNPs). Cyanine 647-labelled fucoidan obtained through aminooxy reaction was used to confirm Fuc-LNPs internalization by immortalized macrophages using confocal microscopy (Figure S4).

**Figure 2.**
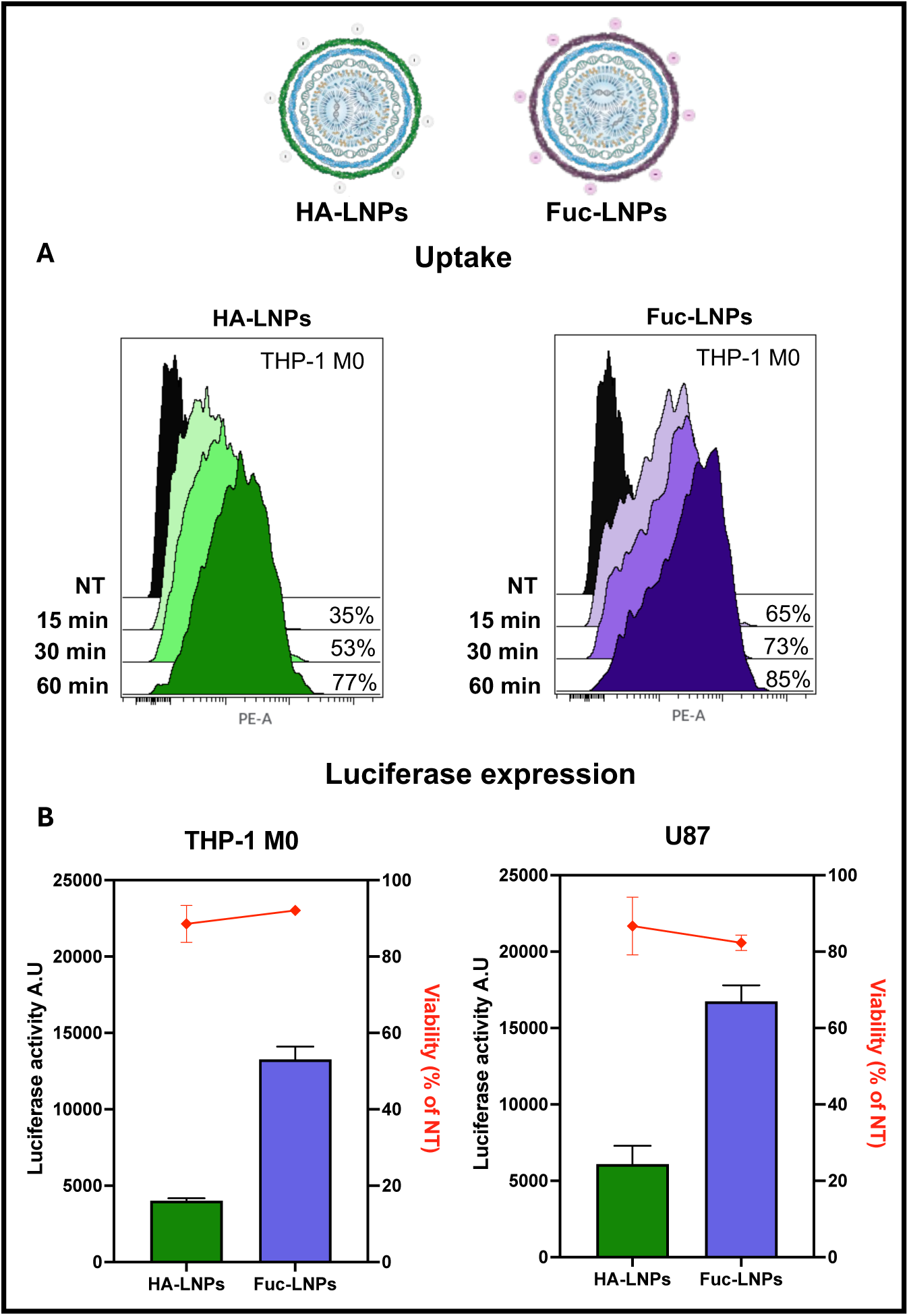
Uptake and mRNA transfection of LbL LNPs in monocytes and macrophages. Flow cytometry uptake of DIR-labelled HA-LNPs and Fuc-LNPs by immortalized (THP-1) derived M0 (top graphs). Cells were incubated with LbL LNPs at equivalent doses of mRNA (50 ng per 50k cells) for 1h after which cells were harvested, washed, and resuspended in FACS buffer for immediate analysis. (B) In vitro transfection in M0 cells (bottom left graph) and U87 glioma cells (bottom right graph) by HA-LNPs or Fuc-LNPs, 48h, 50 ng mRNA Luciferase per well, 50 k cells. Luminescence was measured per protocol (Steady Glow reagent, Promega). Cellular viability was calculated comparing transfected cells to non-treated control via resazurin-based testing.

Subsequently, we wondered how the cellular uptake would be translated into protein expression. We used mRNA encoding the luciferase protein and incubated immortalized M0 with HA-LNPs or Fuc-LNPs. After 48h, we measured luciferase activity and cellular viability. Both LbL LNPs induced little cytotoxicity (red lines, Fig 2B) and Fuc-LNPs promoted a much higher luciferase transfection than HA-LNPs. Measured luminescence for Fuc-LNPs peaked at 15 000 RFU, whereas HA-LNPs remained below 5 000 units (Figure 2B, THP-1 M0 graph).

We wondered if the superior transfection of Fuc-LNPs over HA-LNPs could also be observed in a non-immune cell line. We repeated the experiment, but this time incubating Fuc-LNPs and HA-LNPs with glioma U87 cells, and similar results were observed, where luciferase activity in Fuc-LNPs-treated cells was about 3 times higher than HA-LNPs with no significant difference in induced cytotoxicity for both LbL nanoparticles (Figure 2B, U87 graph).

Having confirmed that fucoidan confers superior colloidal stability to Fuc-LNPs over their hyaluronate counterpart, which is also observed in nanoparticle uptake and mRNA transfection, we started comparing the Fuc-LNPs system with LNPs-PEG (DSPC) formulation in the subsequent experiments. First, if the uptake of Fuc-LNPs could be blocked by free fucoidan, indicating a specific receptor-mediated internalization of particles due to the functionalized outermost layer. To answer this question, we pretreated macrophages with free fucoidan at two different concentrations (50 and 100 μg mL^-1^) before incubation with either Rhodamine-labelled Fuc-LNPs or LNPs-PEG (DSPC) for 2h, when cells were washed and lysed. As expected, both concentrations inhibited the intracellular fluorescence of Fuc-LNPs group, but not the PEGylated LNPs (Figure 3A).

**Figure 3.**
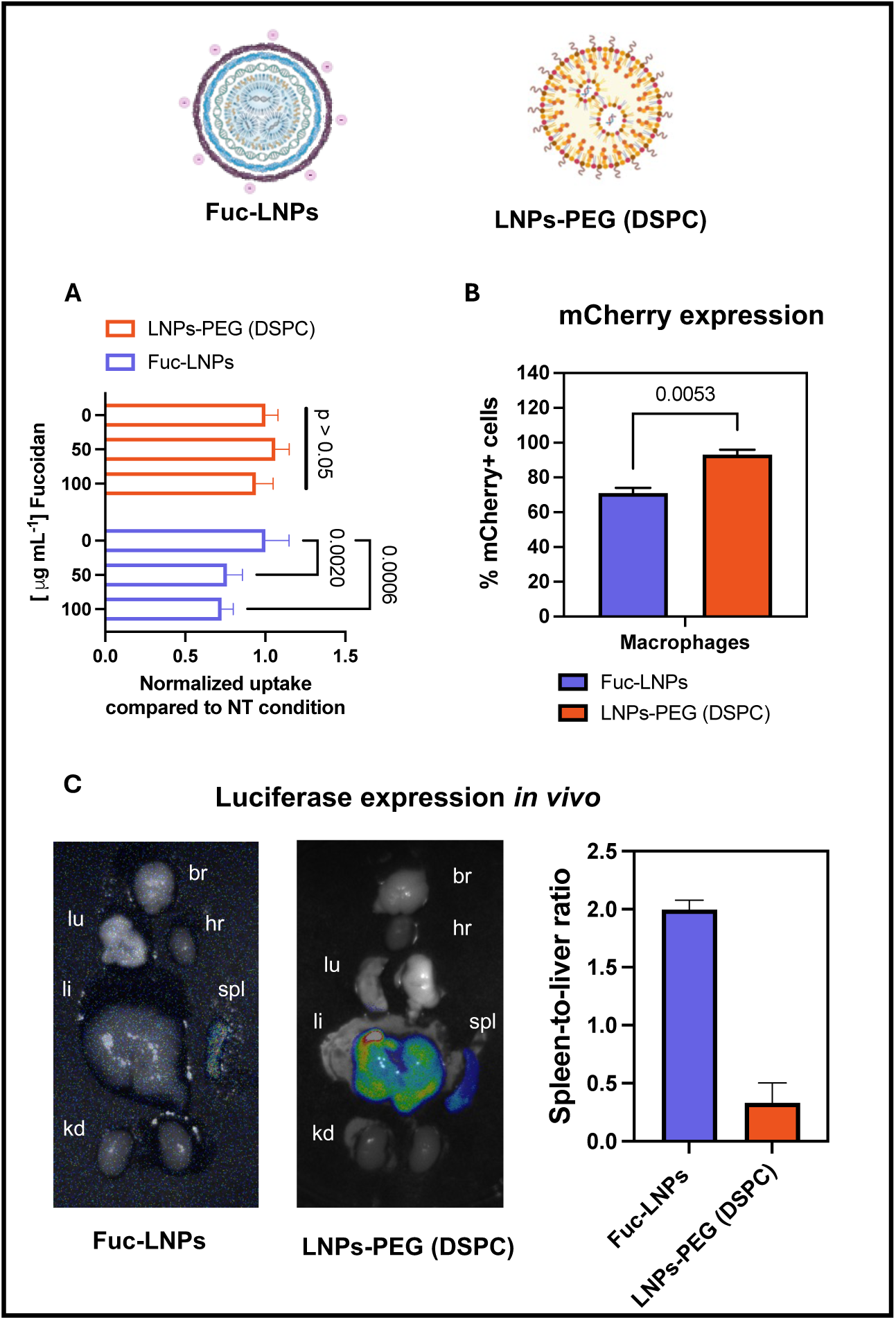
*in vitro* and *in vivo* mRNA transfection of Fuc-LNPs and LNPs-PEG (DSPC) (A) THP-1 monocytes or macrophages transfected with either LNPs-PEG (DSPC) (moderna-like formulation) or Fuc-LNPs loaded with mRNA-mCherry 48h, 200 ng of mRNA per well, 80 k cells (top graph, % of positive cells). (B) In vivo expression of luciferase protein after i.v. injection of LNPs-PEG (DSPC) e or Fuc-LNPs (0.03 mg kg^-1^ of mRNA). Organs were harvested 5h post injection. Organs were identified as follow: Br (brain), hr (heart), lu (lungs), li (liver), spl (spleen), kd (kidneys). Total luminescence (radiance photons/s/cm²/sr) of liver and spleen were used to obtain the spleen-to-liver ratio. Total of 3 animals per condition.

Concerning mRNA transfection in macrophages, Fuc-LNPs induced mCherry expression in about 71% of macrophages, whereas PEGylated LNPs induced a significant higher protein expression (p = 0.0053), reaching approximately 93% of cells (Figure 3B).

We then proceeded to *in vivo* biodistribution experiments. We injected intravenously equivalent mRNA doses for both nanoparticles (0.03 mg of mRNA per kg) and animals were sacrificed 5 h post injection. Results are summarized in Figure 3B. Fuc-LNPs induced almost exclusively spleen transfection whereas luciferase activity was detected in both liver and spleen for LNPs-PEG (DSPC), as expected. The spleen-to-liver ratio for luciferase activity in Fuc-LNPs was 4 times higher than for PEGylated nanoparticles, demonstrating a change in mRNA transfection biodistribution, most likely mediated by the fucoidan-terminated layer-by-layer surface modification of LNPs. Notably, the luciferase activity in Fuc-LNPs treated animals was considerably lower than in PEGylated LNPs.

### Fuc-LNPs have higher tropism for macrophages than PEGylated LNPs in a macrophage-glioma cells co-culture

Based on the high uptake of Fuc-LNPs by macrophages *in vitro*, we wondered how the particles would behave in a glioblastoma-macrophage co-culture. We formulated fluorescently labelled LNPs-PEG (DSPC) and Fuc-LNPs and incubated particles in a co-culture condition containing U87 glioma cells and immortalized human-derived macrophages (HDM) seeded at 1:1 ratio. Cells were incubated with nanoparticles for 1h, fixed and imaged by confocal microscopy (Figure 4). We identified macrophages by their round-shaped morphology (white arrows), as opposed to the epithelial-like morphology of U87 cells. Interestingly, Fuc-LNPs was associated with both macrophages and U87 cells, whereas PEGylated LNPs were captured almost exclusively by U87 cells, with neglectable fluorescence detected in macrophages.

**Figure 4.**
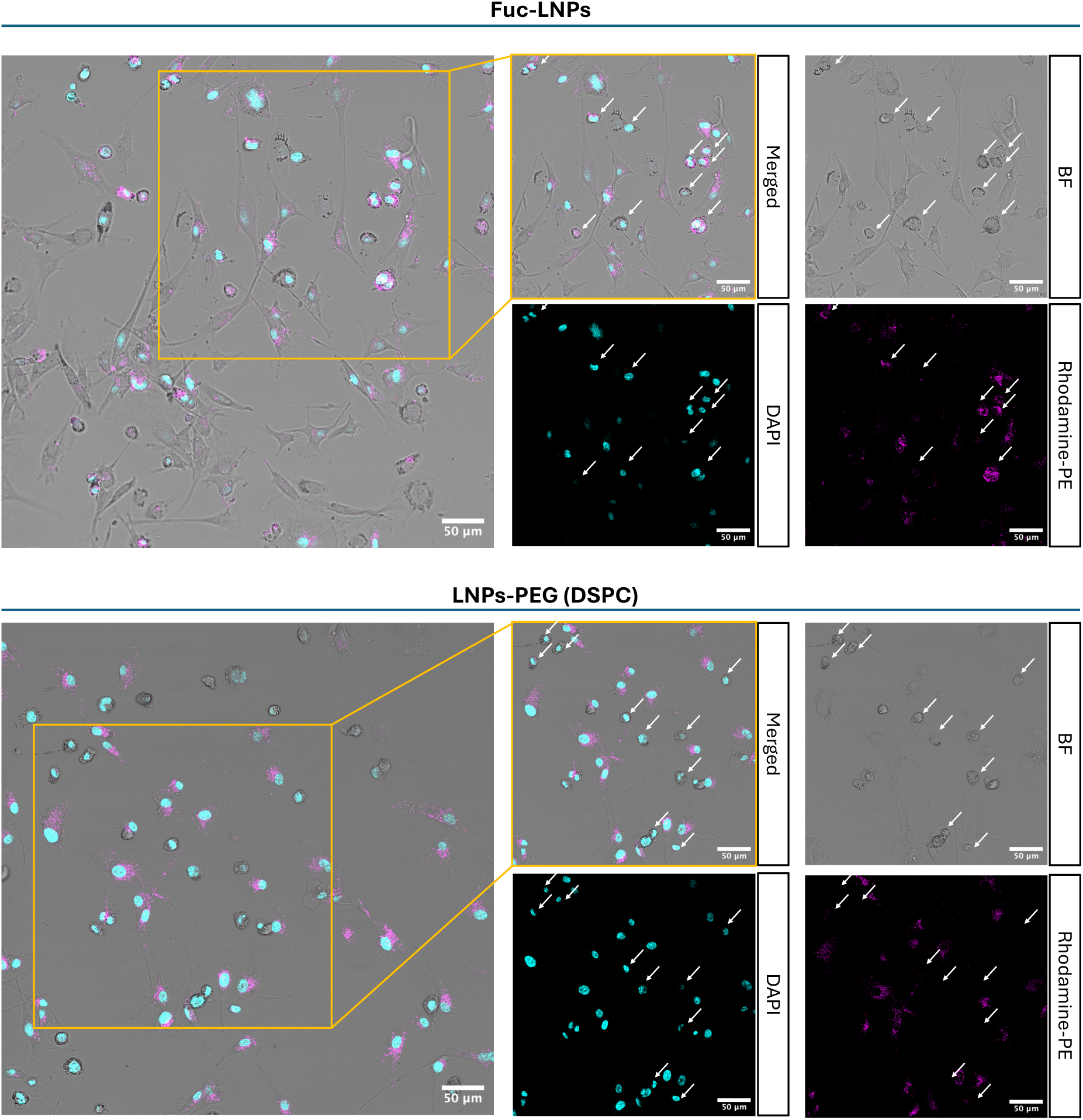
Uptake of Fuc-LNPs and LNPs-PEG (DSPC) in a macrophage/U87 co-culture. THP-1 monocytes were differentiated into macrophages for 48h in the presence of 10 ng mL^-1^ PMA (40 k cells) to yield HDM, after which cells were washed and incubated with 40 k U87 cells overnight. Fuc-LNPs or LNPs-PEG (DSPC) containing Rhodamine-PE lipid (0,1 mol %) were incubated with cells for 2h at equivalent mRNA doses (60 ng of mRNA per well). Cells were washed, fixed, mounted on a coverslip and imaged by confocal microscopy. Signal was colored for interpretation. White arrows indicate round-shaped cells assigned as macrophages.

We then repeated the experiment, but measuring the cellular uptake quantitatively by flow cytometry and discriminating M0 using the CD11b marker. Here we included PEGylated-LNPs formulated with two distinct helper lipids: DSPC (moderna-like formulation, LNPs-PEG DSPC) and DOPE (LNPs-PEG DOPE) for comparative purposes. We started by incubating labelled particles in monoculture conditions and then U87:M0 at 1:1 seeding ratio (Figure 5A).

**Figure 5.**
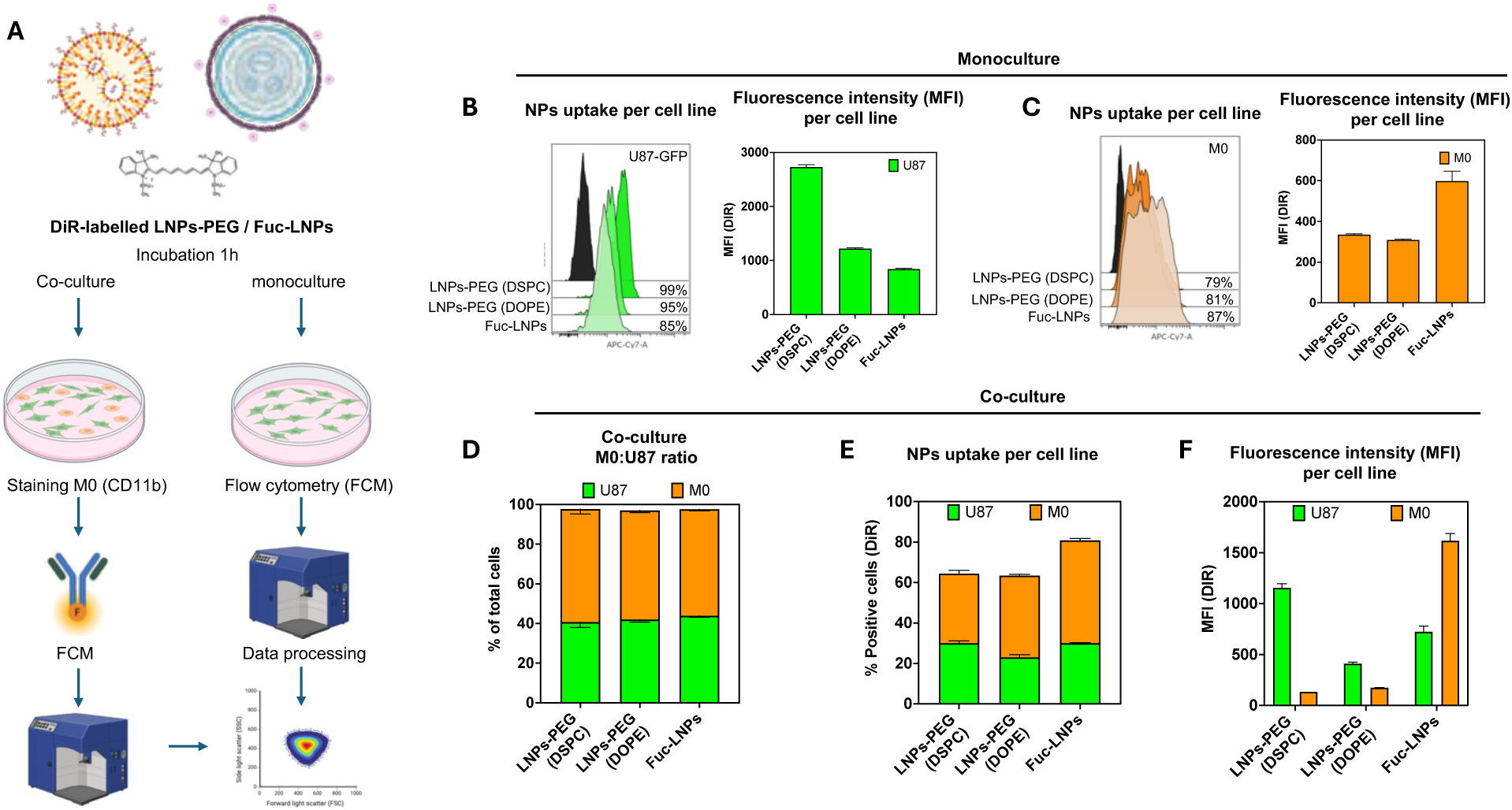
Flow cytometry analysis for the uptake of Fuc-LNPs and PEGylated LNPs in a M0-U87 co-culture condition. (A) Fuc-LNPs and PEGylated LNPs containing DSPC or DOPE were labelled with DiR lipid (0.1 % mol) and incubated in monoculture (B and C) or co-culture conditions (D, E and F, 1:1 cell seeding ratio, 120 k total cells per well), for 1h at equivalent mRNA dose (60 ng of mRNA per well). Cells were washed, harvested, blocked for FcR receptors, and stained for CD11b.

Looking first at the U87-GFP monoculture after 1h incubation with nanoparticles (Figure 5B), we observed the highest uptake from LNPs-PEG containing DSPC (99%), followed by LNPs-PEG (DOPE), 95%, and Fuc-LNPs, 85%. Cells associated with LNPs-PEG (DSPC) also presented the highest median fluorescence intensity (MFI), which was almost three times higher than LNPs-PEG (DOPE) and Fuc-LNPs (Figure 5B). In HDM, however, Fuc-LNPs was associated with 87% of cells, followed by LNPs-PEG (DOPE) with 81% and LNPs-PEG (DSPC), 79%. Fuc-LNPs elicited the highest MFI in macrophages when compared to PEGylated LNPs (Figure 5C).

After observing the behavior of PEGylated LNPs and Fuc-LNPs in isolated U87-GFP and HDM, we proceeded with co-culture conditions. Macrophages-to-glioma cells ratio was consistent in all conditions, with an average of 55% of HDM and 42% U87 cells within the single cell gated population (Figure 5D). Regarding uptake, LNPs-PEG (DSPC) associated with HDM and U87 to similar extents, 30% and 34%, respectively (Figure 5E), but MFI associated with HDM was considerably low (Figure 5F). In LNPs-PEG (DOPE) condition, we observed a shift towards macrophage-targeting over U87 cells, with 40% association in HDM and 23% in glioma cells (Figure 5E). However, for both PEGylated conditions, the MFI in U87 cells was higher than in macrophages (Figure 5F). Still, the overall fluorescence for the cellular association of LNPs-PEG (DOPE) was lower than LNPs-PEG (DSPC).

For Fuc-LNPs, more than 50% of macrophages were associated with the LbL system, whereas this value dropped to 30% for U87 cells (Figure 5E). MFI for Fuc-LNPs associated with HDM was two-fold higher than the one observed for U87-GFP cells, a clear inversion in the pattern observed for PEGylated LNPs (Figure 5F). As a consequence, Fuc-LNPs elicited a superior macrophage targeting and associated-cellular fluorescence across all nanoparticles tested in this co-culture condition. Altogether, both flow cytometry and confocal microscopy indicate that the presence of fucoidan in the outermost layer of LbL LNPs confer macrophage tropism to lipid nanoparticles.

### Fuc-LNPs efficiently generate CAR-M in vitro

We next investigated the ability of Fuc-LNPs in producing CAR-macrophages (CAR-M) *in vitro.* We carried out transfection in both immortalized and primary human-derived macrophages. First, in primary cells, we isolated CD14+ monocytes from peripheral blood from healthy donors, which were then differentiated into macrophages following a 6-days protocol under macrophage colony stimulating factor (M-CSF). At the end of the differentiation period, macrophages were transfected with either PEG-LNPs (DPSC) or Fuc-LNPs loaded with mRNA encoding luciferase or the CD19-targeted CAR (CAR@CD19) (Figure 6).

**Figure 6.**
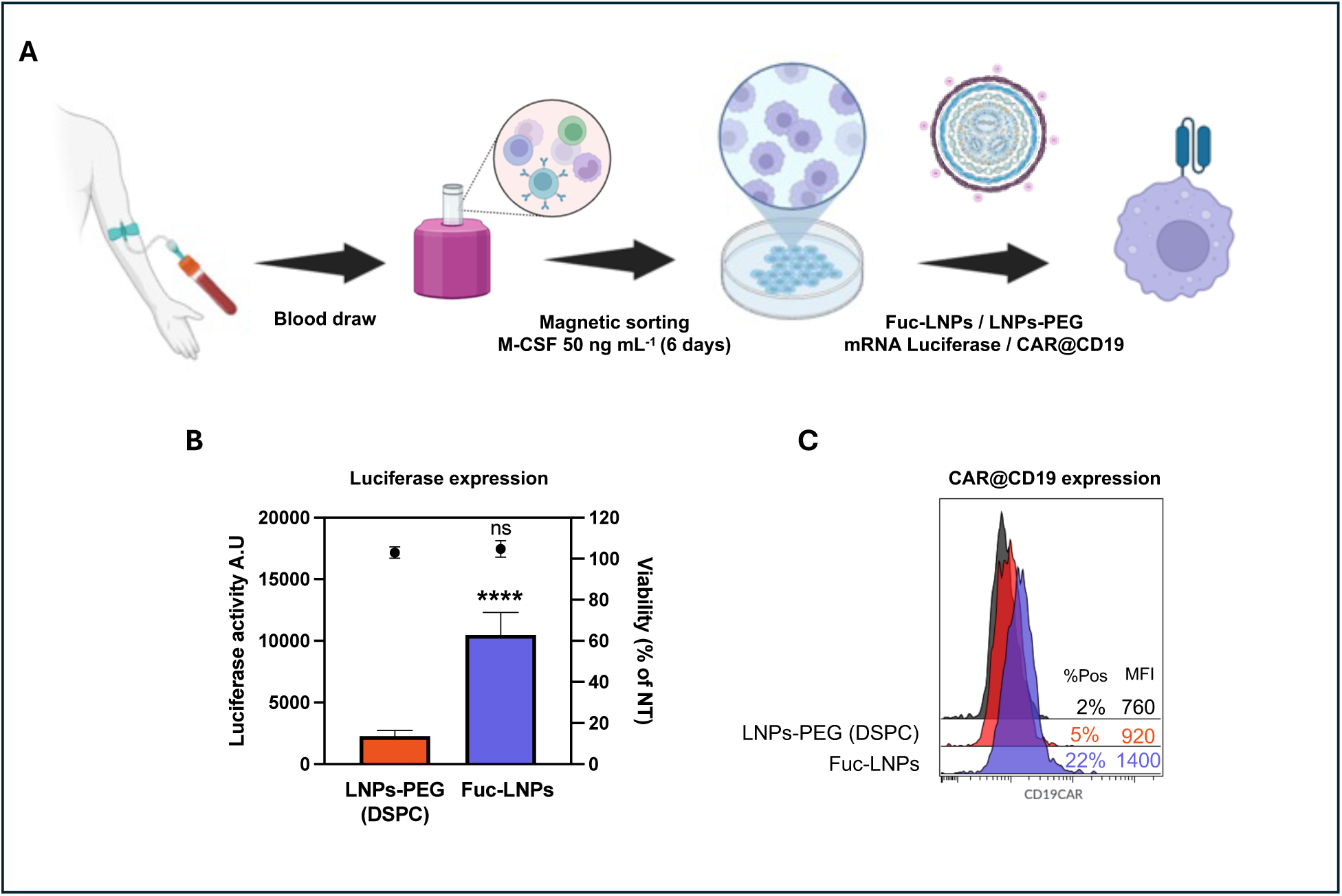
Transfection of primary human-derived macrophages. (A) Monocytes were isolated from healthy donors and differentiated into macrophages under stimulus of M-CSF for 6 days, after which cells were transfected with Fuc-LNPs or LNPs-PEG (DSPC). For mLuciferase experiments (B), cells were transfected with 42 ng of mRNA per well (60 k cells); for CAR@CD19 experiments (C), primary HDM were transfected with 200 ng of mRNA per well (250 k cells). After 48h, cells were collected for flow cytometry or processed for luciferase activity / cellular viability per protocol.

Following the isolation and differentiation of human-derived primary macrophages (Figure 6A), we started by evaluating transfection efficiency by transfecting mRNA encoding the luciferase protein. 48h after transfection, we measured both cellular viability and luciferase activity. No toxicity was induced by either formulation, but Fuc-LNPs induced a superior luciferase transfection when compared to LNPS-PEG (DSPC) as indicated by a 4-fold higher enzymatic activity than the one observed for their PEGylated counterpart (Figure 6B). Finally, we asked if the superior mRNA transfection for Fuc-LNPs in primary macrophages would be observed for mRNA encoding the CD19-targeting CAR. Similarly to mRNA reporter experiments, we transfected primary cells with Fuc-LNPs or PEGylated LNPs for 48h, when the cells were harvested, blocked, and stained with APC-tagged recombinant CD19 protein to recognize CAR-expressing macrophages (Figure 6C). Fuc-LNPs was more effective than LNPs-PEG (DSPC) in generating CAR-Macrophages *ex vivo* with a positivity rate for the surface expression of CAR-CD19 of 22% for Fuc-LNPs (MFI of 1400) versus 5% for PEGylated LNPs (MFI of 920). In immortalized HDM, the CAR@19 positivity induced by Fuc-LNPs reached 33% (Figure S6A), therefore we chose to test the functionality CAR-M against CD19-expresing cells using THP-1 derived macrophages.

To evaluate the cytotoxic effect of CAR-Macrophages, we incubated CAR@CD19 (Fuc-LNPs) or untransfected macrophages (UNT-M0) with Raji-GFP/Luciferase cells at 1:1 effector to target cells ratio (E:T) for 48h. We monitored the GFP intensity by live tracking microscopy using the Incucyte platform (Figure 7A) and we enzymatically measured the luminescence activity from Raji cells at the end of the incubation period (Figure 7B). CAR@CD19 induced significantly lower levels of luminescence than UNT-M0, indicating a higher cytotoxicity of transfected macrophages against Raji cells. Indeed, we observed a decrease in the normalized Raji-GFP signal by live tracking microscopy when these cells were incubated with CAR@CD19 versus UNT-M0. Curiously to observe if the same effect would be reproduced in an adherent cell line, we induced the expression CD19 protein in U87-GFP/Luc cells (78% positivity, 48h post transfection, Figure S6B) and co-incubated with CAR-M. Again, we observed a decreased tumor growth by live tracking microscopy (Figure 7C), a lower luciferase activity (Figure 7D) at the end of 48h co-incubation. Furthermore, at the 24h time point, we detected an increased expression of TNF-α release in the CAR-M + U87CD19 condition when compared to UNT-M0 co-incubation (Figure S6C). To visualize the interaction between macrophages and U87CD19-GFP/Luc cells, we performed confocal microscopy at the 24 h time point in a parallel, illustrative experiment (Figure 7E). In both conditions, macrophages established physical contacts with tumor cells. In the CAR-M co-cultures, small GFP-positive puncta were occasionally observed within IBA1⁺ macrophages, indicating limited transfer of tumor-derived material (Figure S5). However, we did not observe complete engulfment of intact U87 cells. The SIRPα/CD47 axis is a strong inhibitory axis present in solid tumors that prevents the phagocytic activity of myeloid cells. Indeed, both macrophages (untransfected and CAR-M) and U87-GFP/Luc cells expressed high levels of SIRPα (Figure S6D) and CD47 (Figure S6E) proteins, respectively.

**Figure 7.**
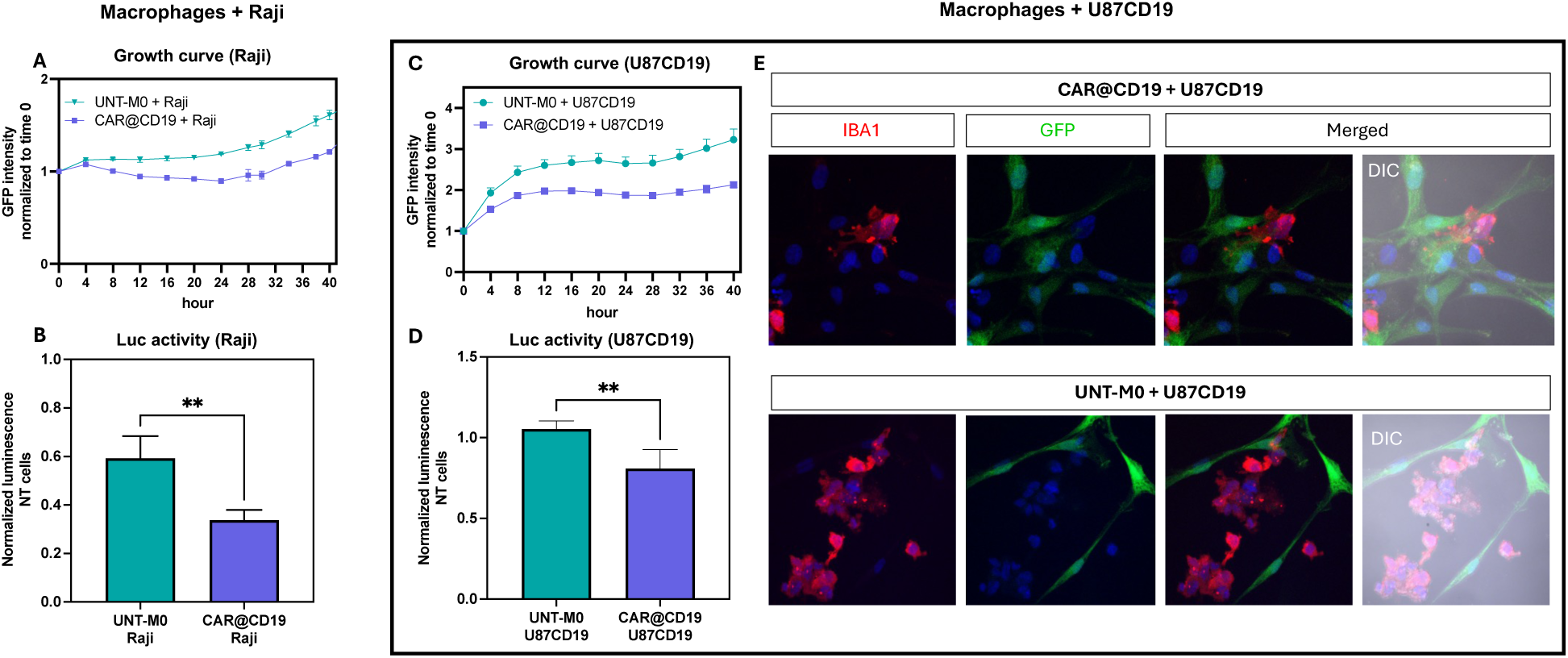
Transfection of primary human-derived macrophages. (A) CAR@CD19 cells were generated from THP-1 derived M0 (200 ng mRNA per well) and incubated with either Raji-GFP/Luc or U87CD19-GFP/Luc cells for 48h, where GFP signal was tracked by live imaging (A and C) and luciferase measured at the experiment endpoint (B and D). At 24h time point, macrophages (UNT-M0 and CAR@CD19) co-incubated with U87CD19-GFP/Luc were fixed, intracellular staining for IBA1, and imaged by confocal microscopy (E).

Altogether, the results summarized in this investigation demonstrate that layer-by-layer lipid nanoparticles containing fucoidan as the outermost layer possess distinguished physicochemical behavior when compared to their unPEGylated lipidic template and the hyaluronate layer-by-layer counterpart. Additionally, fucoidan provided macrophage tropism to LbL LNPs, which was not observed for in PEGylated LNPs formulations. Finally, Fuc-LNPs successfully engineered macrophage *ex vivo* inducing the surface expression of CD19@CAR to a superior extent than PEGylated formulation in primary macrophages.

## Discussion

In this study, we report on the functionalization of lipid nanoparticles using the layer-by-layer technique for efficient macrophage engineering through RNA modulation. By choosing fucoidan as outermost polyelectrolytes in the 3-layered system, we bestowed colloidal stability to unPEGylated mRNA-loaded LNPs when diluted in PBS. Additionally, Fuc-LNPs sustained lyophilization in the presence of cryoprotectant (Figures 1C, 1D). Notably, after lyophilization, Fuc-LNPs maintained size, PDI and negative surface charge upon resuspension in PBS which was not observed with PEGylated LNPs. Additionally, resuspended Fuc-LNPs induced similar levels of transfection than fresh formulation using the *in vitro* model HEK 293T cells (Figure S1C). Interestingly, without cryoprotectant, Fuc-LNPs increase in size after lyophilization, although to a lesser extent than LNPs-PEG in similar conditions (data not shown), underscoring the importance of the modular selection of polyelectrolytes, notably the outermost layer, during the LbL modification of LNPs. Indeed, controlling ionic strength is an important parameter when addressing colloidal stability of electrostatic assembled LbL systems since it directly impacts polyelectrolytes spatial disposition and rearrangement and their biological outcome (29). To the best of our knowledge, this is the first comprehensive study on the colloidal stability of LbL-assembled nanoparticles in PBS.

This finding immediately drove us to elucidate the mechanism whereby the sulfated polysaccharide fucoidan, but not the carboxylated sodium hyaluronate, protects the LbL LNPs from aggregation in such high ionic strength solution. We observed that the lipidic core is still accessible to small molecules sensing either the ionizable lipid protonation state (Figure S3A) or the RNA templates (Figure S3B). Nabar and coworkers (30) investigated the accessibility of nanoparticles to TNS and reported a significant decreased pK_a_^A^ of monolayered LNPs when compared to the unlayered counterparts. Additionally, researchers observed a buffering zone in the TNS assay caused by the two ionizable species in the system. Although we expected a similar behavior in the LbL LNPs developed here, these phenomena were not observed, except for a higher apparent pK_a_ of Fuc-LNPs than PEGylated unlayered nanoparticles (LNPs-PEG DSPC and LNPs-PEG DOPE).

The elucidation of the LNPs’ internal structure by SAXS also suggest a unique feature of LbL LNPs as they presented lower q-values than PEGylated LNPs and the unlayered unPEGylated template, indicating an increased internal d-spacing.

Nano flow cytometry revealed that not all Fuc-LNPs contained mRNA-Cy5 (39.4% mRNA-loaded v. 60.6 % mRNA-empty Fuc-LNPS, Figure 1B, Figure S2A). It is fair to assume this value would also be found for PEGylated-LNPs. This assumption is corroborated by similar q-values for unPEGylated LNPs and LNPs-PEG indicated by SAXS measurements (Figure S3C). However, during the LbL process, both mRNA-loaded and empty core templates are complexed with an additional stiffer siRNA layer that is successfully found in more than 97% of nanoparticles (NanoFCM, Figure S2A), fundamentally changing the structural organization of vesicles in the layer-by-layer condition, probably explaining the left-shifted q-value for Fuc-LNPs, which present a higher RNA loading per particle than the PEGylated LNPs.

The left-shift in Braggs peaks for PEGylated LNPs as one raises the pH is a feature commonly reported (31) and was also observed in this study (Figure S1C). However, for Fuc-LNPs, the diffraction patterns were completely abrogated upon dilution in PBS. It remains to be investigated if the signal loss was a consequence of the decrease in concentration caused by the buffer exchange to PBS and the following purification step for SAXS measurement.

Fucoidan has been exploited in macrophage-targeting therapies. In fact, fucose-rich, sulphate-containing polysaccharides extracted from brown seaweed are reported to interact with macrophages via several receptors, as 4-1BB (32), Scavenger Receptors class A (33, 34) , and TLR4 (35). Although most studies on fucoidan have concentrated on the polysaccharide’s effect on macrophage polarization and anti-tumoral properties (36), here we leveraged the sulfated polyelectrolyte’s affinity towards macrophages to demonstrate efficient mRNA delivery to murine and human-derived macrophages. In immortalized human macrophages, Fuc-LNPs induced a higher transfection efficiency when compared to hyaluronan-decorated LbL LNPs (Figure 2B). Moreover, as a P-selectin ligand (27), Fuc-LNPs also induced strong mRNA transfection in U87 glioma cell lines (Figure 2B). P-selectin is overexpressed in endothelial cells and many cancer types (27, 37), including brain tumors where fucoidan functionalized nanoparticles demonstrated selectin-mediated blood-brain barrier crossing and significant tumor accumulation (38, 39).

In the co-culture experiment, Fuc-LNPs surpassed both PEGylated LNPs in macrophage targeting, although DOPE-containing LNPs-PEG performed slightly better than the DSPC counterpart, aligning with evidence that softer nanoparticles showed increased uptake in macrophages (40). Considering the Fuc-LNPs formulated herein contain DOPE as helper lipid, and the layer-by-layer assembly seem to minimally contribute to the elastic moduli of nanoparticles (41), which comes predominantly from the lipidic core template, one may assume that both mechanical properties of LbL LNPs and fucoidan-driven targeting may be contributing towards a macrophage tropism in the co-culture experiment.

Consideration should be given to the experiments with primary macrophages. Macrophages are highly specialized in recognizing foreign RNA and, upon immune activation, mount an immune response shutting down translational machinery, thus negatively affecting gene transfection. In the case of CAR expression, a new barrier in transfection efficiency is the translocation of the newly synthesized protein to the surface membrane of macrophages. The PEGylated LNPs induced a minimal surface expression of CAR@CD19, whereas Fuc-LNPs maintained a significant potency in mRNA transfection (Figure 6C), albeit lower than observed for immortalized macrophages (Figure S6A). Additionally, here we chose to use a mRNA encoding the CAR@CD19 bearing classical intracellular activation domains (4-1BB / CD3ζ) and N1-methylpseudouridine base modification, which both can be optimized for enhanced macrophage activation and expression (42, 43). Finally, using THP-1 derived macrophages as a source to generate CAR@CD19 Macrophages, we observed that Fuc-LNPs-transfected cells induced ablation of CD19-expressing cells at 1:1 E:T ratio (Figure 7). Importantly, the high expression of “don’t eat me” signal may have helped U87CD19 cells evading myeloid cells phagocytosis (44). Indeed, targeting the SIRPα-CD47 axis improve tumor cells clearance by CAR-Macrophages (45–47).

## Conclusion

Here we reported on the development of a novel fucoidan-decorated layer-by-layer LNPs containing mRNA for the modulation of macrophages. We demonstrate that the outermost layer impacted the stability of nanoparticles, providing a PEG-free strategy to stabilize mRNA-loaded LNPs in iso-osmolar medium (PBS). In addition, Fuc-LNPs sustained lyophilization, maintaining physiochemical properties and functionality, assessed by mRNA transfection *in vitro*. The layer-by-layer strategy allows double RNA loading, opening up the possibility of delivering a core-protected mRNA and a layered RNAi for a dual-therapeutic effect. To conclude, here we presented a novel strategy to engineer human macrophages using layer-by-layer-modified mRNA-loaded LNPs expanding the toolbox for CAR therapies. Further experiments will focus on the development of CAR-macrophages also addressing their polarization and validating the cellular therapy in solid tumors *in vitro* and *in vivo*.

## Acknowledgements

PH acknowledge the support from CHU Sainte-Justine (Fonds d’innovation thérapeutique). VPG acknowledges the training funds from the Vanier Canada Graduate scholarship operated by the Canadian Institutes of Health Research (CIHR 475599). This work is based upon research conducted at the Structural Biology Platform of the Université de Montréal (RRID:SCR_022303). The Platform was funded and is currently supported by the Canadian Foundation for Innovation award #30574. We acknowledge Anton Paar for providing access to the Litesizer DLS 700 instrument used in this study.

## CRediT authorship contribution statement

**VPG:** Conceptualization, Methodology, Formal analysis, Investigation, Writing - original draft. **HT:** Methodology, Supervision, Writing - review & editing. **AF and JS:** Investigation, formal analysis. **SO:** Investigation, Writing - review & editing **NB:** Resources, Supervision. **MCBLN**: Writing - review & editing **XB:** Conceptualization, Writing - review & editing, Funding acquisition, Supervision. **P H:** Conceptualization, Methodology, Resources, Writing - review & editing, Funding acquisition, Supervision.

