## Supporting information for "Fucoidan-coated Layer-by-layer Lipid Nanoparticles for the generation of CAR-Macrophages"

[f] Laboratório de Nanotecnologia, Biotecnologia e Cultura de Células, Centro Acadêmico de Vitória, Universidade Federal de Pernambuco (CAV/UFPE), 55608-680 Recife, Brazil

[g] Child Health and Human Development Axis, Research Institute of the McGill University Health Centre, Montréal, Québec, Canada; Department of Human Genetics, McGill University, Montréal, Québec, Canada.

[h] Department of Pediatrics, Université de Montréal, Montréal, QC, H3C 3J7, Canada

**Keywords:** Layer-by-layer Lipid Nanoparticles, RNA delivery, CAR-Macrophages, Immunotherapy, Glioblastoma

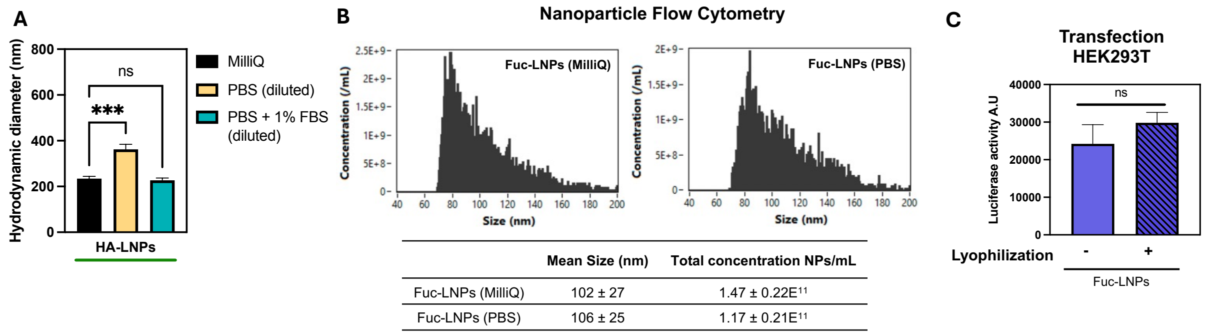


**Figure S1**.(A) Hydrodynamic diameter expressed as Z-Average. Hyaluronan-decorated LNPs were diluted in pure water, PBS, or PBS supplemented with 1 % FBS. Particles were incubated for 10 minutes, then measured using the LiteSizer DLS 700. HA-LNPs diluted in pure water v. dilution in PBS, **** ($p < 0.0001$); HA-LNPs diluted in pure water v. dilution in PBS + 1 % FBS, ns, $p > 0.05$. (B) Fucoidan LbL LNPs (Fuc-LNPs) were diluted in pure water or PBS and submitted to nanoparticle flow cytometry (NanoFCM) using the scattering signal to measure particle size distribution and concentration. (C) Lyophilized Fuc-LNPs were resuspended in PBS to the initial volume and incubated with HEK-293T cells (50 ng of mRNA Luciferase per 50 k cells/well) for 48h. Luciferase activity was measured using Steady-Glo reagent. No statistical difference was found between samples.

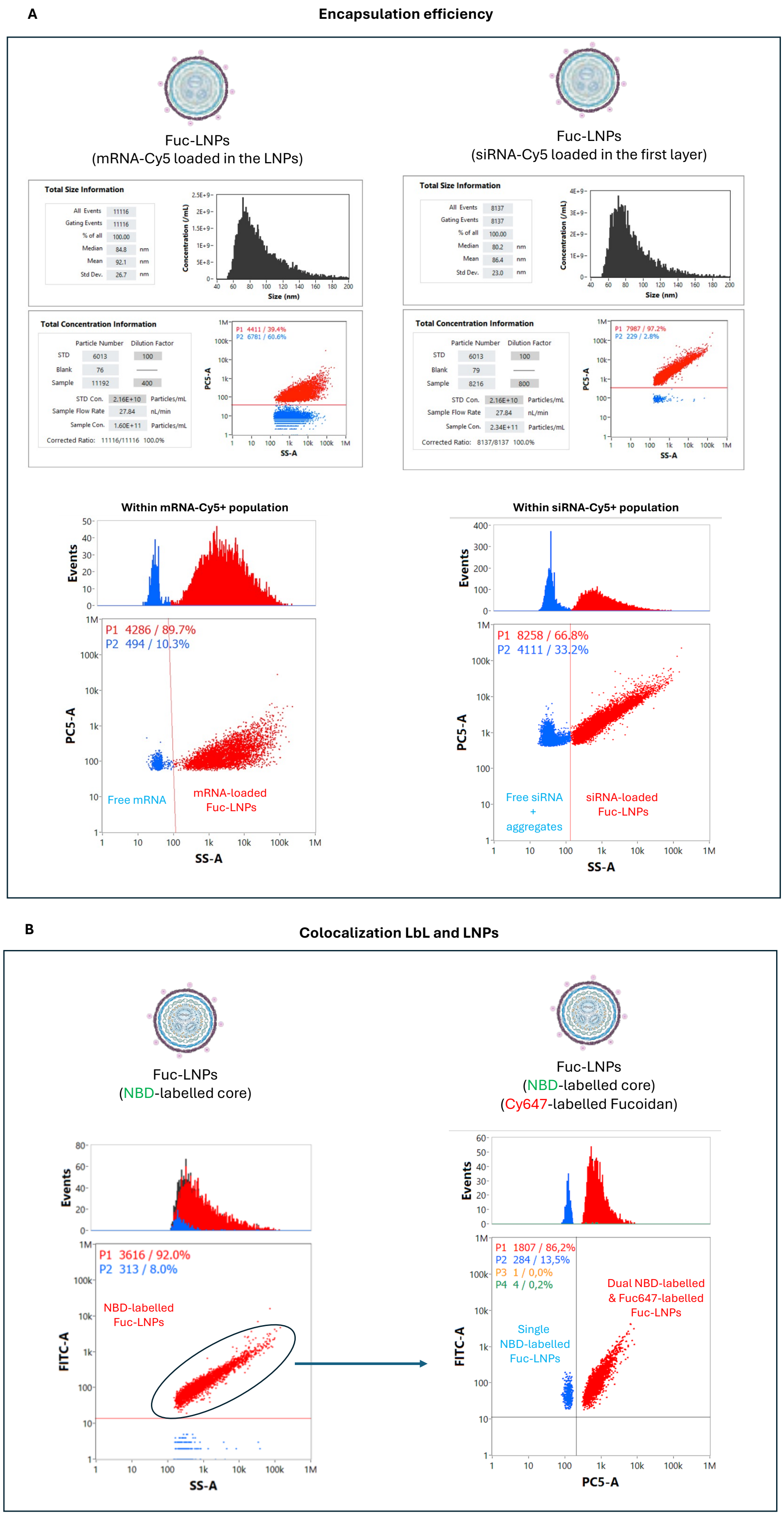
A
Encapsulation efficiency
Fuc-LNPs
(mRNA-Cy5 loaded in the LNPs)
Fuc-LNPs
(siRNA-Cy5 loaded in the first layer)
Total Size Information
All Events 11116
Gating Events 11116
% of all 100.00
Median 84.8 nm
Mean 92.1 nm
Std Dev. 26.7 nm
Concentration (/mL)
Size (nm)
Total Size Information
All Events 8137
Gating Events 8137
% of all 100.00
Median 80.2 nm
Mean 86.4 nm
Std Dev. 23.0 nm
Concentration (/mL)
Size (nm)
Total Concentration Information
Particle Number Dilution Factor
STD 6013 100
Blank 76
Sample 11192 400
STD Con. 2.16E+10 Particles/mL
Sample Flow Rate 27.84 nL/min
Sample Con. 1.60E+11 Particles/mL
Corrected Ratio: 11116/11116 100.0%
P1 4411 / 39.4%
P2 6781 / 60.6%
PC5-A
SS-A
Total Concentration Information
Particle Number Dilution Factor
STD 6013 100
Blank 79
Sample 8216 800
STD Con. 2.16E+10 Particles/mL
Sample Flow Rate 27.84 nL/min
Sample Con. 2.34E+11 Particles/mL
Corrected Ratio: 8137/8137 100.0%
P1 7987 / 97.2%
P2 229 / 2.8%
PC5-A
SS-A
Within mRNA-Cy5+ population
Events
P1 4286 / 89.7%
P2 494 / 10.3%
PC5-A
Free mRNA
mRNA-loaded
Fuc-LNPs
SS-A
Within siRNA-Cy5+ population
Events
P1 8258 / 66.8%
P2 4111 / 33.2%
PC5-A
Free siRNA
+
aggregates
siRNA-loaded
Fuc-LNPs
SS-A
B
Colocalization LbL and LNPs
Fuc-LNPs
(NBD-labelled core)
Fuc-LNPs
(NBD-labelled core)
(Cy647-labelled Fucoidan)
Events
P1 3616 / 92.0%
P2 313 / 8.0%
NBD-labelled
Fuc-LNPs
FITC-A
SS-A
Events
P1 1807 / 86,2%
P2 284 / 13,5%
P3 1 / 0,0%
P4 4 / 0,2%
Dual NBD-labelled
& Fuc647-labelled
Fuc-LNPs
Single
NBD-labelled
Fuc-LNPs
FITC-A
PC5-A

**Figure S2**. (A) Fuc-LNPs were formulated with either mRNA-Cy5 or siRNA-Cy5, diluted in Pure water and analysed using NanoFCM. (B) Fuc-LNPs were formulated with NBD-PE (0.1 % mol), left graph, or NBD-PE and Cyanine647-conjugated Fucoidan, right graph, and submitted to NanoFCM.

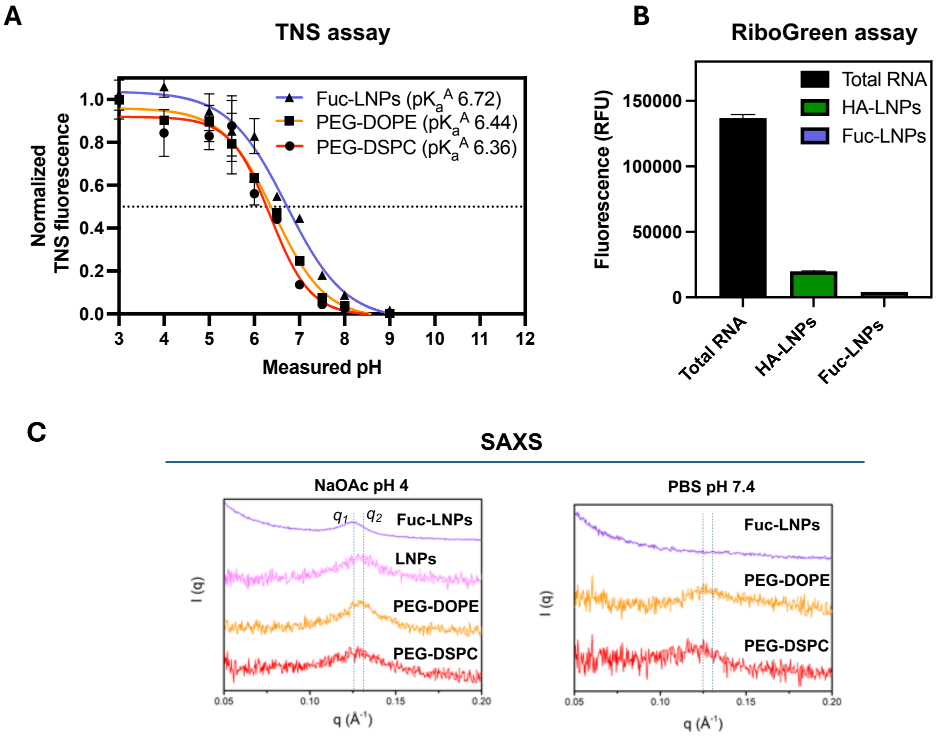


**Figure S3**. (A) TNS assay of Fuc-LNPs and PEGylated-LNPs containing either DSPC or DOPE as helper lipids. (B) The intercalating agent, RiboGreen, was used to determine the accessibility of HA-LNPs and Fuc-LNPs. Total fluorescence was compared to an RNA solution containing mRNA and siRNA at equivalent molar concentration than LbL LNPs. (C) SAXS measurements for LbL LNPs containing fucoidan (Fuc-LNPs), unPEGylated LNPs core (LNPs) or PEGylated LNPs (PEG-DOPE and PEG-DSPC). Particles were diluted in either pH 4 buffer (sodium acetate, 10 mM, pH 4) or PBS prior concentration using Amicon filters (100 kDa).

A

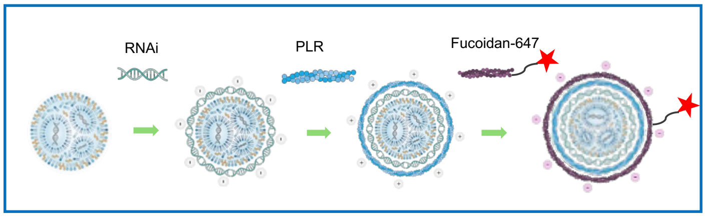


B

| | Mean Size (nm) | Total concentration NPs/mL | Size (Zave) nm | PDI | ζ-potential (mV) |
|---|---|---|---|---|---|
| Fuc647-LNPs (MilliQ) | 98 ± 22 | 1.60 ± 0.22E$^{11}$ | 182 ± 3 | 0.215 ± 0.06 | -24 ± 1 |
| Fuc647-LNPs (PBS) | 111 ± 31 | 0.85 ± 0.22E$^{11}$ | 201 ± 2 | 0.168 ± 0.01 | -27 ± 1 |

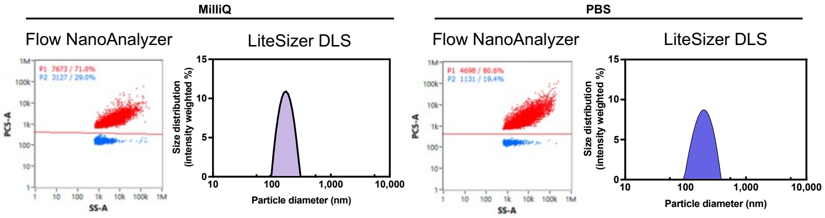


C

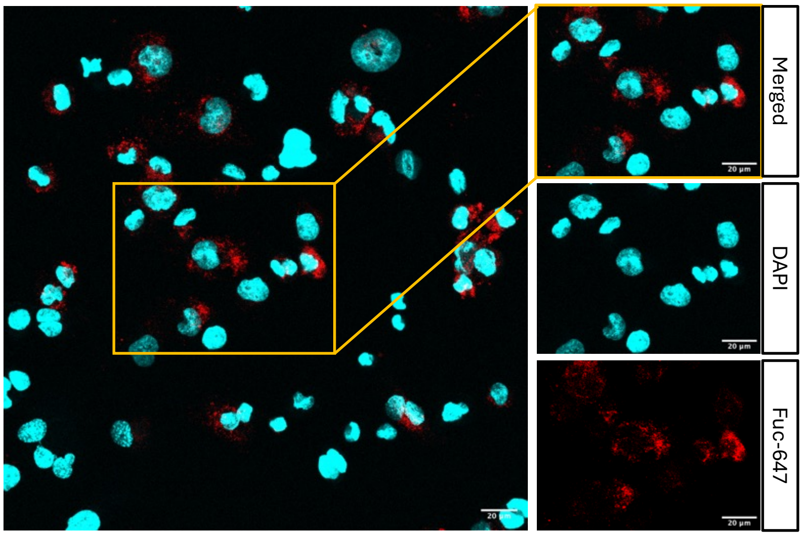


**Figure S4**. (A) Schematic representation for the synthesis of Cyanine647-conjugated Fucoidan as the outermost layer of Fuc-LNPs. (B) Physicochemical characterization using NanoFCM of Fuc-LNPs diluted in Pure water or PBS. (C) Confocal microscopy of THP-1 derived macrophages incubated with Cyanine647-conjugated Fucoidan decorated LNPs.

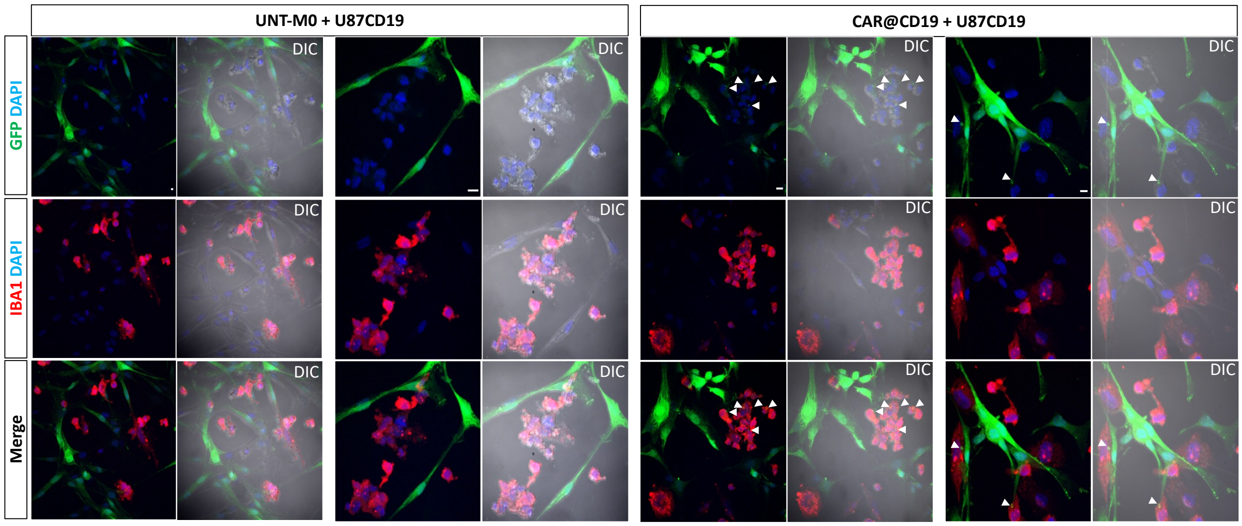


**Figure S5.** Spatial organization and cell-cell interactions within the co-culture at the 24-hour time point. Representative confocal images of U87-GFP tumor cells (green) co-cultured with macrophages stained for IBA1 (red). Nuclei are labeled with DAPI (blue), and DIC is shown to visualize overall cell morphology. White arrows highlight small GFP-positive particles located within the cytoplasm of IBA1$^{+}$ macrophages. Scale bar = 10um

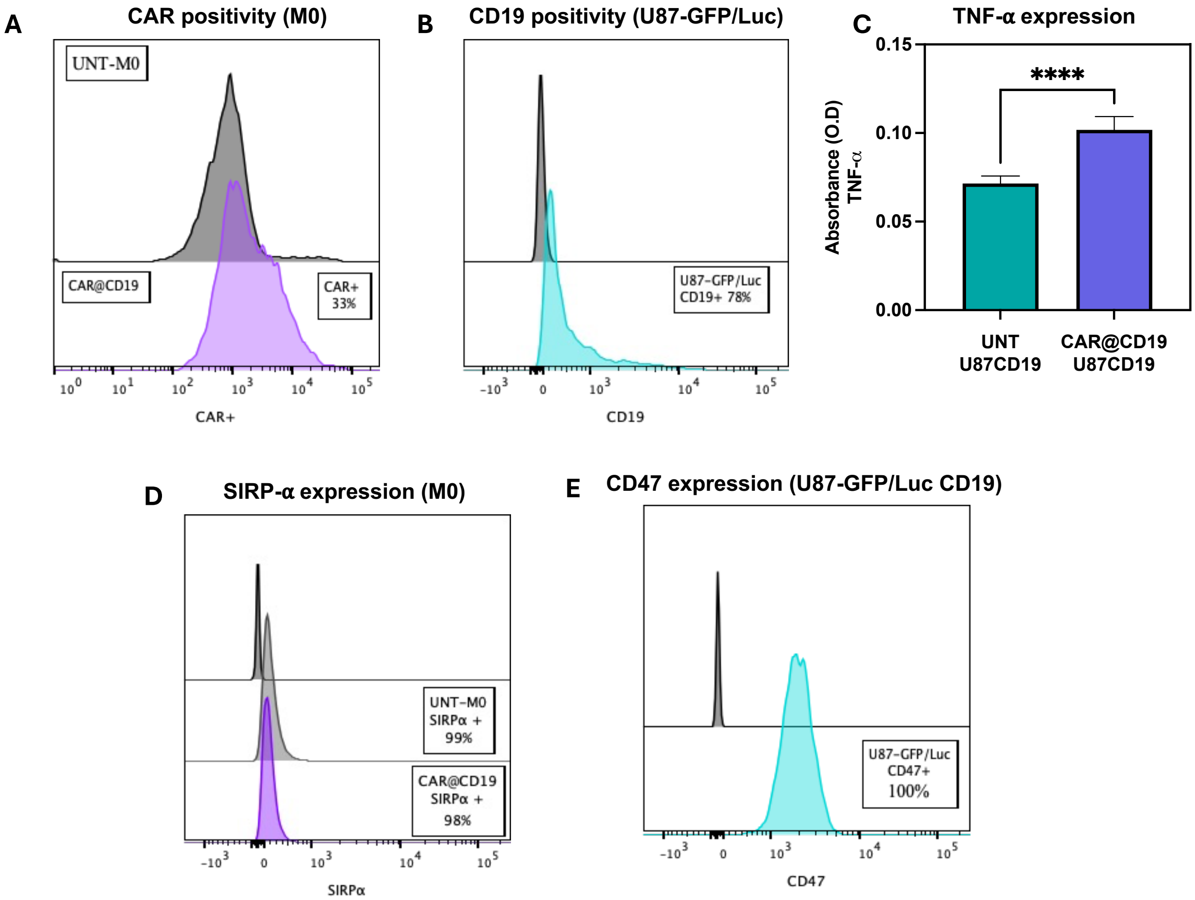


**Figure S6.** (A) Detection of CAR@CD19 in THP-1 derived macrophages transfected with Fuc-LNPs (200 ng / mRNA / 200 k cells / 48h). (B) U87-GFP/Luc cells were transfected with Lipofectamine 3000 per manufacturer's protocol and surface expression of CD19 was detected 48h post transfection. (C) TNF-α was detected in the supernatant of untransfected or CAR@CD19 macrophages incubated (24h) with U87-GFP/Luc expressing CD19. (D) and (E) surface expression of SIRPα and CD47 surface proteins in macrophages and U87-GFP/Luc cells, respectively, by flow cytometry, 24h after seeding.
